# Patterns of polymorphism at the self-incompatibility locus in 1,083 *Arabidopsis thaliana* genomes

**DOI:** 10.1101/098228

**Authors:** Takashi Tsuchimatsu, Pauline M. Goubet, Sophie Gallina, Anne-Catherine Holl, Isabelle Fobis-Loisy, Héléne Bergés, William Marande, Elisa Prat, Dazhe Meng, Quan Long, Alexander Platzer, Magnus Nordborg, Xavier Vekemans, Vincent Castric

**Affiliations:** Univ. Lille, CNRS, UMR 8198 - Evo-Eco-Paleo, F-59000 Lille, France; Gregor Mendel Institute, Austrian Academy of Sciences, Vienna Biocenter, Vienna, Austria; Department of Biology, Chiba University, Yayoi-cho 1-33, Inage, Chiba 263-8522, Japan; Reproduction et Développement des Plantes, Institut Fédératif de Recherche 128, Centre National de la Recherche Scientifique, Institut National de la Recherche Agronomique, Université Claude Bernard Lyon I, Ecole Normale Supérieure de Lyon, Lyon, France; Centre National des Ressources Génomiques Végétales, INRA UPR 1, Castanet-Tolosan, France; Dept. of Biochemistry and Molecular Biology & Cumming School of Medicine. University of Calgary.

## Abstract

Although the transition to selfing in the model plant *Arabidopsis thaliana* involved the loss of the self-incompatibility (SI) system, it clearly did not occur due to the fixation of a single inactivating mutation at the locus determining the specificities of SI (the S-locus). At least three groups of divergent haplotypes (haplogroups), corresponding to ancient functional S-alleles, have been maintained at this locus, and extensive functional studies have shown that all three carry distinct inactivating mutations. However, the historical process of loss of SI is not well understood, in particular its relation with the last glaciation. Here, we took advantage of recently published genomic re-sequencing data in 1,083 *Arabidopsis thaliana* accessions that we combined with BAC sequencing to obtain polymorphism information for the whole S-locus region at a species-wide scale. The accessions differed by several major rearrangements including large deletions and inter-haplogroup recombinations, forming a set of haplogroups that are widely distributed throughout the native range and largely overlap geographically. ‘Relict’ *A*. *thaliana* accessions that directly derive from glacial refugia are polymorphic at the S-locus, suggesting that the three haplogroups were already present when glacial refugia from the last Ice Age became isolated. Inter-haplogroup recombinant haplotypes were highly frequent, and detailed analysis of recombination breakpoints suggested multiple independent origins. These findings suggest that the complete loss of SI in *A*. *thaliana* involved independent self-compatible mutants that arose prior to the last Ice Age, and experienced further rearrangements during post-glacial colonization.

## Introduction

The transition from outcrossing to self-fertilization (selfing) is one of the most frequent evolutionary processes in flowering plants (Stebbins 1974, Barrett 2002, Igic et al. 2008, Shimizu and Tsuchimatsu 2015). Although selfing may lead to inbreeding depression, it has advantages *via* reproductive assurance (Darwin 1876), and also has automatic transmission advantage (Fisher 1941). In obligate outcrossers, self-fertilization is often avoided by post-pollination genetic self-incompatibility (SI) (Igic et al. 2008, De Nettancourt 2001, Takayama and Isogai 2005); thus the transition to selfing necessarily involves the loss of the SI system.

Given its evolutionary prevalence, the loss of SI has been a major research focus for plant evolutionary biologists (Igic et al. 2008, Charlesworth and Charlesworth 1979, Uyenoyama et al. 2001, Busch and Schoen 2008, Gervais et al. 2014). SI systems generally consist of genes determining self-recognition specificities and other genes involved in signaling pathways that lead to pollen rejection when pollen and pistils express cognate specificities (de Nettancourt 2001, Takayama and Isogai 2005). The male and female specificity genes are generally tightly linked at a single genetic locus (the S-locus) with multiple highly diverged alleles segregating (called S-alleles, S-haplotypes or S-haplogroups, de Nettancourt 2001). In the Brassicaceae, where the SI system has been extensively studied, the S-locus receptor kinase (SRK) and the S-locus cysteine-rich proteins (SCR; also called S-locus protein 11, SP11) determine the female and male specificities, respectively. A few other genes involved in the signaling pathway downstream of *SRK* have also been identified, such as the M-locus protein kinase (MLPK), the ARM-repeat Containing1 (ARC1), and the exocyst subunit Exo70A1 (Stone et al. 2003, Murase et al. 2004, Samuel et al. 2009, Indriolo et al. 2012; but see Kitashiba et al. 2011).

*Arabidopsis thaliana* has been studied extensively as a model for the loss of SI (Kusaba et al. 2001, Nasrallah et al. 2002, Nasrallah et al. 2004, Sherman-Broyles et al.2007, Liu et al. 2007, Tang et al. 2007, Shimizu et al. 2008, Tsuchimatsu et al. 2010, Dwyer et al. 2013). *A*. *thaliana* is self-compatible and predominantly selfing (Abbott and Gomes 1989, Platt et al. 2010, Bomblies et al. 2010), while closely related species such as *A*. *lyrata* and *A. halleri* are mainly obligate outcrossers and posess the ancestral Brassicaceae SI system (Mable et al. 2003, Llaurens et al. 2008). Analysis of the *SRK* gene identified three highly divergent but non-functional S-haplogroups in *A*. *thaliana*: A, B and C (Shimizu et al. 2008). Functional sequences very similar to each of these three S-haplogroups were found in the outcrossing relatives *A*. *lyrata* and *A*. *halleri*, demonstrating that the *A*. *thaliana* alleles derive from distinct ancestral S-alleles (Bechsgaard et al. 2006).

A major challenge in the study of the loss of SI has been determining the sequence of events, and in particular distinguishing primary inactivating mutations from secondary decay after initial loss of function (Igic et al. 2008, Busch and Schoen 2008, Tsuchimatsu et al. 2010, Boggs et al. 2009). Sequence analysis of the S-locus region identified several gene-disruptive mutations in both *SCR* and *SRK* (Kusaba et al. 2001, Nasrallah et al. 2004, Tang et al. 2007, Shimizu et al. 2008, Tsuchimatsu et al. 2010). When functional copies of these genes from *A*. *lyrata* were transformed into *A. thaliana,* SI reactions were restored in several accessions (Nasrallah et al. 2002, Liu et al. 2007, Dwyer et al. 2013, Boggs et al.2009), suggesting that the mutations in the S-locus genes may have been responsible for the loss of SI in *A. thaliana,* although the contribution of other loci such as *ARC1* and *PUB8* remains controversial (Indriolo et al. 2012, Indriolo et al. 2014, Nasrallah and Nasrallah 2014, Goring et al. 2014). One study described a 213-bp inversion of the *SCR* gene in haplogroup A and reported that restoring the ancestral arrangement was sufficient to restore functional SI (Tsuchimatsu et al. 2010). This result suggested that degradation of *SCR* was one of the primary mutations responsible for the loss of SI. However, the inversion is not fixed in the species: it is restricted to accessions carrying haplogroup A (Tsuchimatsu et al. 2010). In parallel, several structural rearrangements of the S-locus have been reported, including inter-haplogroup recombination between haplogroups A and C as well as large deletions encompassing S-locus genes (Sherman Broyles et al. 2007). While these rearrangements could obviously disrupt the gene functions and therefore may have contributed to the evolutionary loss of SI, they too may simply represent secondary decay after the loss of SI.

To chart the historical degradation of the S-locus, it is essential to have a detailed view of polymorphism of the whole region from geographically widely distributed samples. However, this is challenging because the S-locus region has extremely high nucleotide diversity, structural rearrangements and repetitive sequences that make traditional sequence analysis impracticable (Guo et al. 2011, Goubet et al. 2012). Previous studies in *A*. *thaliana* were based on small numbers of short fragments obtained by PCR-based genotyping, Sanger sequencing, or DNA gel blotting for a limited set of accessions (Nasrallah et al. 2004, Sherman Broyles et al. 2007, Tang et al. 2007, Shimizu et al. 2008, Boggs et al. 2009).

Recently, whole-genome information has become publicly available for more than one thousand worldwide accessions in *A*. *thaliana* (Cao et al. 2011, Gan et al. 2011, Long et al. 2013, The 1001 Genomes Consortium 2016). We took advantage of these data to explore species-wide polymorphism of the whole S-locus region. We disentangled the complicated mutational history of this region by making use of the raw read sequences from natural accessions. First, as a reference for the short-read mapping, we obtained the full S-locus sequence of haplogroup C, to complement the existing sequences for A and B (Tang et al. 2007, AGI 2000; but see also Dwyer et al. 2013) and we functionally characterized the self-incompatibility genes. Second, we mapped the short reads of each accession to the reference sequences of each of the A, B, and C S-haplogroups. Based on these data, we consider how ancestral S-haplotypes have lost functionality independently. *A*. *thaliana* survived the last glacial age as a set of isolated lineages in separate glacial refugia (The 1001 Genomes Consortium 2016). While the extent of postglacial recolonization of most lineages remained limited, descendants of one of the refugia drastically expanded and invaded most of the European continent, admixing with local lineages. Interestingly, several accessions show no evidence of admixture with the invaders, i.e. they appear to be ‘relicts’ of their respective glacial refugia (The 1001 Genomes Consortium 2016, Lee et al. 2017). By including these accessions, we show that selfing must have originated before the last glacial period. We find a high frequency of haplotypes originating from inter-haplogroup recombination, and show that these were formed by many independent recombination events in the population that successfully recolonized most of Eurasia.

## Results

### Polymorphism and divergence among copies of haplogroup C

Only a partial sequence was available for the S-locus of haplogroup C (from accession Lz-0, Dwyer et al. 2013), so we first constructed a bacterial artificial chromosome (BAC) library and obtained the complete sequence of the S-locus region from the accession Ita-0, which also carries haplogroup C (Shimizu et al. 2008). We recovered an 11-kb sequence fragment with 97% overall identity to the previous data from haplogroup C sequence, including in the intergenic regions (from ATREP20 to ARK3; Fig. 1). However, there are major structural differences between the two sequences, including a large 25-kb fragment present only in Ita-0 (from TAT1_ATH to ARNOLDYlc, Fig. 1). Notably, Ita-0 also lacks two peculiar features of the S-locus sequence of Lz-0, namely the insertion of a truncated duplicated copy of the flanking gene *ARK3* (called *ΔARK3)* as well as the five fragmented duplicates of *SRK* described in Dwyer et al. (2013), demonstrating that these structural changes are not fixed within haplogroup C. Overall, this direct comparison demonstrates that several types of structural polymorphisms, including large indels and local rearrangements involving gene fragments, exist within haplogroup C. To polarize the observed polymorphisms, we analyzed the full sequence of the S-locus region for a copy of the functional ortholog of haplogroup C in *A*. *halleri* (Ah36), also obtained from a dedicated BAC library (Bechsgaard et al. 2006, Durand et al. 2014). The coding sequence of *SCR* and *SRK* showed high conservation between Ah36 and haplogroup C (Fig 2A and 2B), but most intergenic regions within the S-locus showed too little sequence conservation for reliable polarization of the polymorphisms segregating in *A*. *thaliana* (Fig 2A).

**Figure 1.**
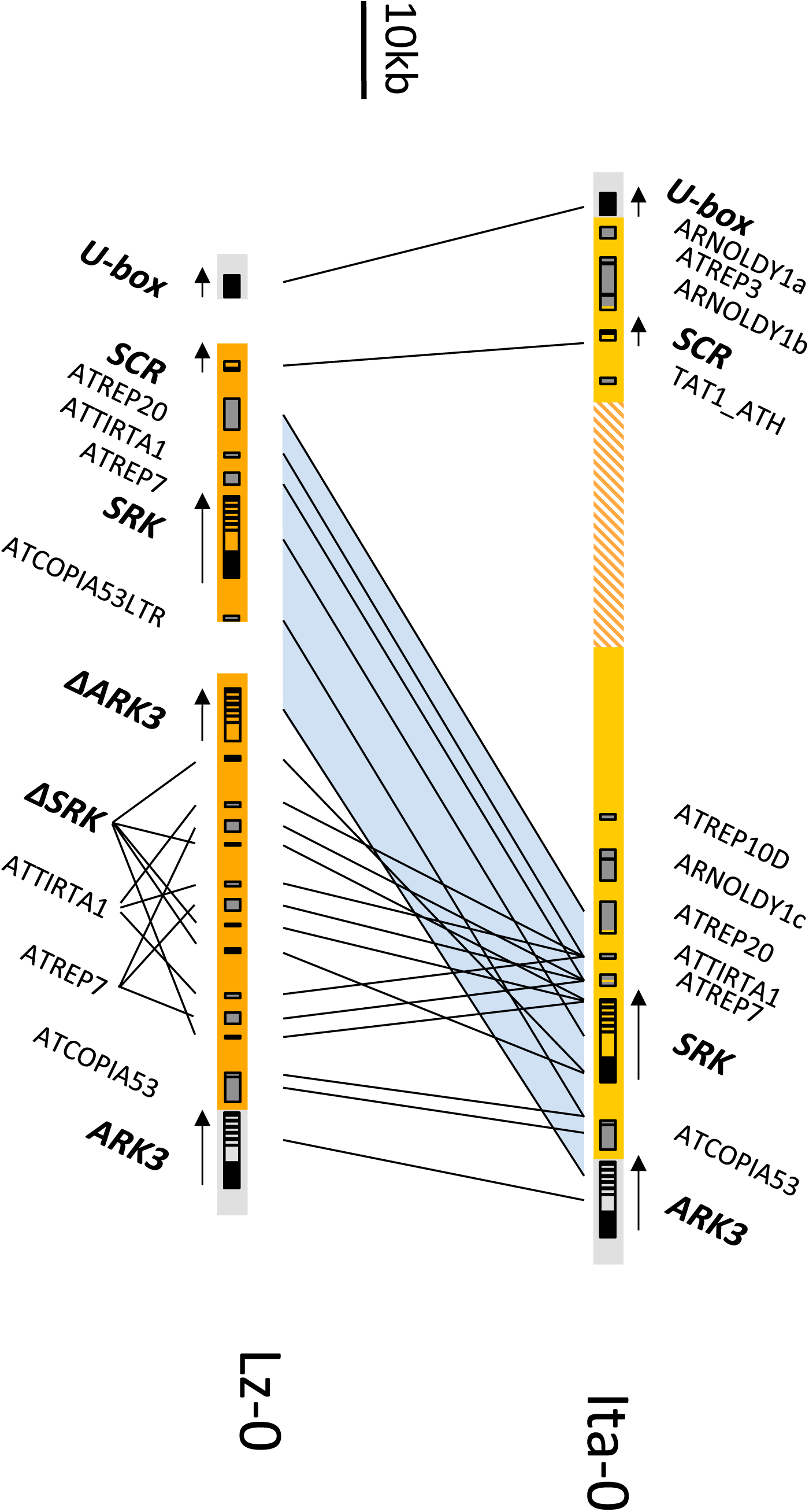
The S-locus region of haplogroup C from Ita-0 compared with Lz-0. Annotations were performed independently and lines join identical annotations in the two sequences.

### Lack of *SCR* expression associated with a large deletion in the promoter in haplogroup C

Dwyer et al. (2013) showed that *SRK* is expressed normally in Lz-0 but has inactivating mutations in its coding sequence, while the coding sequence of *SCR* is functional but poorly expressed. We confirmed a similar pattern in Ita-0: we found substantial expression of *SRK* in stigmatic tissues but no detectable *SCR* transcripts in anthers (Fig. 2C). Based on our complete sequences of the Ita-0 and Ah36 S-locus, we further observed high conservation of the promoter region of *SRK* over about 440bp (Fig. 2B), whereas the 5’ flanking region of *SCR* was extremely diverged from that of the functional AhSCR36 (Fig. 2B), possibly accounting for the reduction of *SCR* expression. *SRK* amino-acid sequence comparison shows that Ita-0 is an outgroup to all other members of haplogroup C (Fig. S1), suggesting that the rearrangements (as well as any other putative inactivating mutations) occurred before the divergence between Ita-0 and other accessions of haplogroup C.

**Figure 2.**
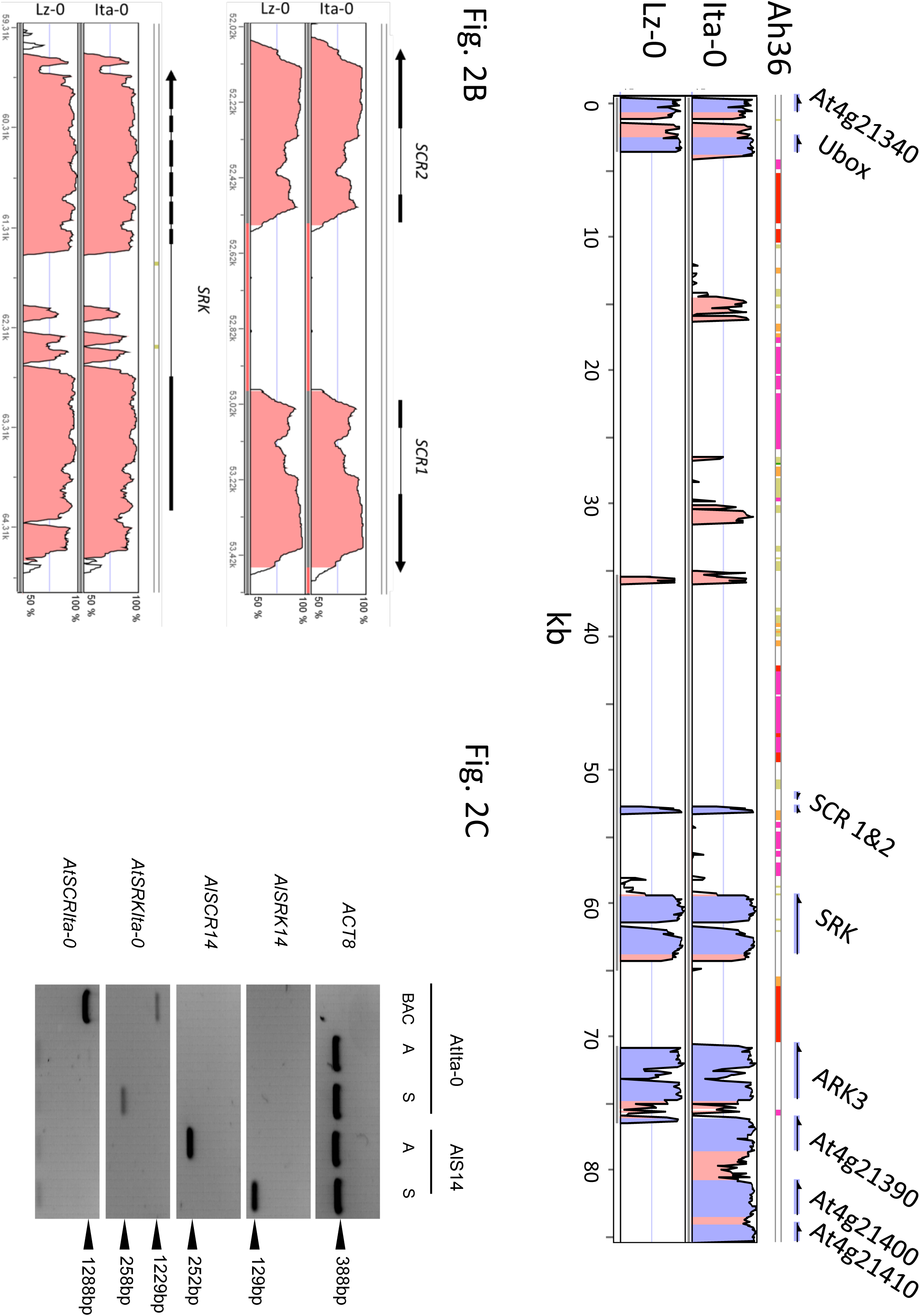
Polymorphism and divergence within haplogroup C. **A.** Vista plot comparing the S-locus region of Ah36 from *Arabidopsis halleri* (used as a reference) with those of Lz-0 and Ita-0 accessions. The grey bars below the Lz-0 sequence represent the three non-contiguous fragments obtained by Dwyer et al. (2012) **B.** Zoom on the *SCR* and *SRK* regions. Ah36 is used as a reference. **C.** Expression analysis of *SCR* and *SRK* in anthers (A) and stigmas (S) of the Ita-0 accession. *SCR14* and *SRK14* from *A*. *lyrata* extracts are used as positive controls. Expression level of *AtSCRIta-0* in anther was very low, as we did not detect any PCR product even after 35 PCR cycles.

### Identification of diverse sequence types by mapping to multiple references

We then took advantage of recently published genomic re-sequencing data in 1,083 *A*. *thaliana* accessions from throughout the species range (The 1001 Genomes Consortium 2016) to document the species-wide molecular diversity of the S-locus genomic region. The high density of transposable elements which characterizes the S-locus region (Goubet et al. 2012) prevents accurate *de novo* assembly of the region. Furthermore because S-locus haplogroups are highly diverged from one another, alignment to the *A*. *thaliana* reference genome is impracticable. Hence, we aligned the raw short reads to available references for each of the three haplogroups: haplogroup A from Col-0 (AGI 2000), haplogroup B from Cvi-0 (Tang et al. 2007) and haplogroup C from Ita-0 (described above).

We first generated coverage plots against each haplogroup reference for every accession (Fig 3). Strikingly, the vast majority of accessions could be categorized into a small number of groups with similar patterns of quantitative coverage to the three reference sequences (Fig. 4). Closer examination of the coverage plots allowed us to identify nine distinctive contiguous fragments of the references (Cov_1, 3 and 4 on haplogroup A and Cov1, 2, 3, 4, 6 and 7 on haplogroup C, Fig. S2) that varied qualitatively among accessions and showed a clear bimodal distribution of coverage (Fig. S3). We scored these fragments as either present or absent, leading us to distinguish a total of twelve sets of accessions, plus one singleton sequence, based on the composition of the S-locus (Fig. 4). These sets included three different groups of “pure A” (A1, A2, A3), one limited group of only two “pure B” accessions and three groups of “pure C” accessions (C1, C2, C3). In addition, five groups of accessions showed substantial coverage on both A and C references, and presumably correspond to recombinant accessions (see below). Sanger sequencing confirmed that the border of the deletion of the Cov-2 fragment (distinguishing A2 and R2 haplotypes from the rest of A-carrying accessions) was the same as that previously identified in C24 (Sherman Broyles et al. 2007). A single accession (Db-1) did not appear to match any of the three reference sequences (Fig. 4), and may represent either a fourth ancestral haplogroup (which would be present at extremely low frequency), or a rare complete deletion of the S-locus specific to this accession.

**Figure 3.**
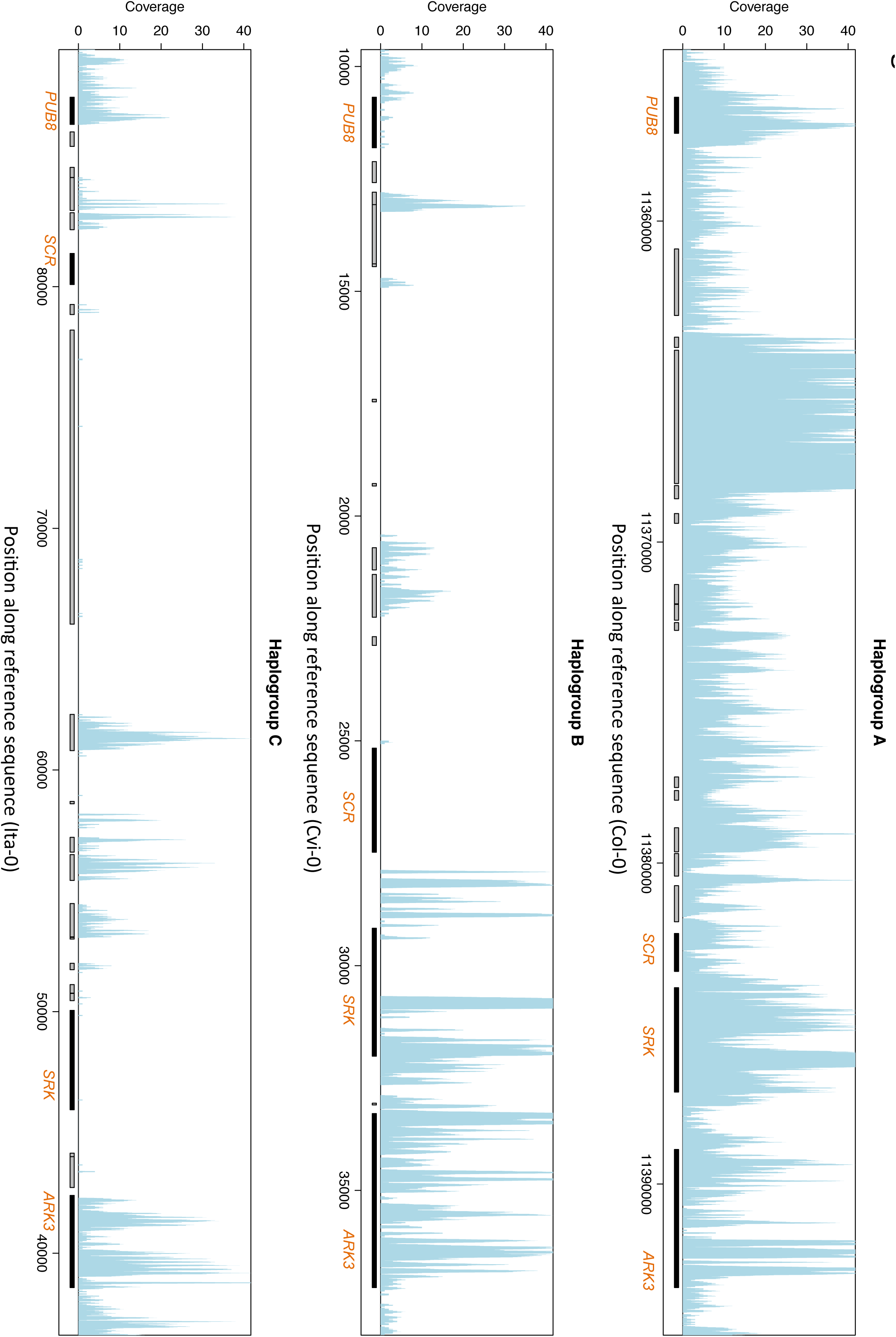
Example coverage plots for one of the 1083 accessions (accession Moran-1, haplotype A1). Coverage represents the number of sequencing reads for each position of the three reference sequences from haplogroups A, B and C. Regions between two flanking genes *(ARK3* and *PUB8)* are shown. Transposable elements (TE) and genes are represented by grey and black horizontal bars, respectively. Note that even though this accession is assigned as haplotype A1, some coverage is apparent on the *SRK* gene of the B reference, an artifact due to some sequence similarity between A and B *SRK* alleles (Bechsgaard et al. 2006). Note the important coverage on the TE sequences due to their repetitive nature, but which is not informative with regard to haplotype assignment.

**Figure 4.**
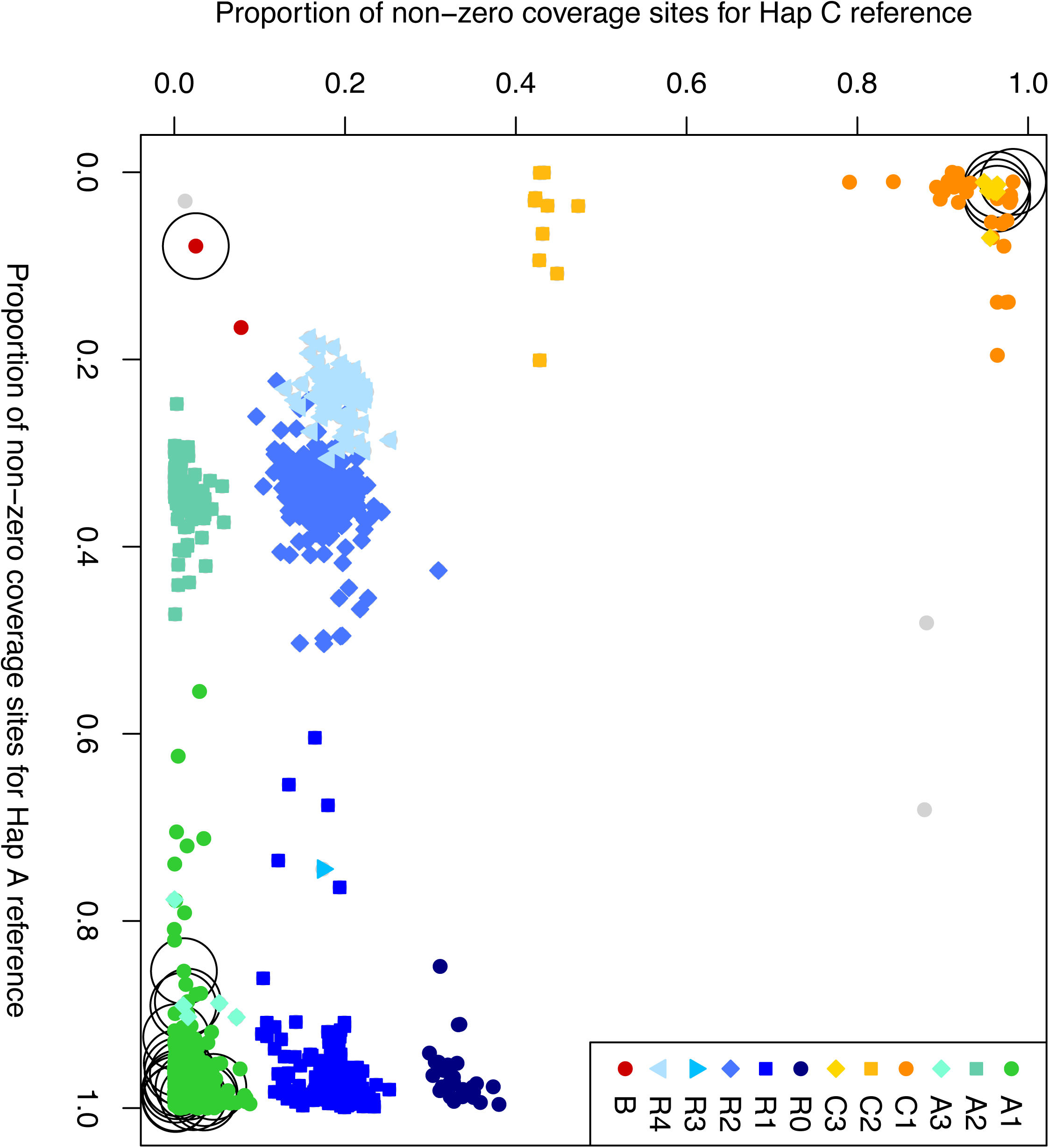
Proportion of nucleotide sites with non-zero coverage on the A and C references. Accessions are color-coded according to their pattern of presence/absence of the fragments defined in Fig. S2. ‘Relict’ accessions are circled with a black circle in the background. Coverage on the B reference was consistently low except for two accessions (Cvi-0 and Can-0) and is not shown here.

### Geographical distribution of S-haplogroups

The data resulting from the classification described above provide a species-level distribution of S-locus genotypes — the first in any species (Fig. 5). Globally, haplogroup A is common (43.21%), while haplogroup C is rare (5.38%), and haplogroup B is extremely rare (two accessions, 0.18%). Surprisingly, the remaining accessions carry recombinant haplotypes (51.5 %, see below). Haplogroup A was widely distributed across Europe but was particularly common in the east. In contrast, haplogroup B was only found in the two accessions in which it had already been reported, i.e. in the Canary and Cape Verde Islands. Haplogroup C was more common in western Europe, and it always occurred together with with haplogroup A and recombinant haplotypes. Altogether, both A and C are found across Europe, with a tendency for higher frequency of A in the east and C in the west (Fig. S4).

**Figure 5.**
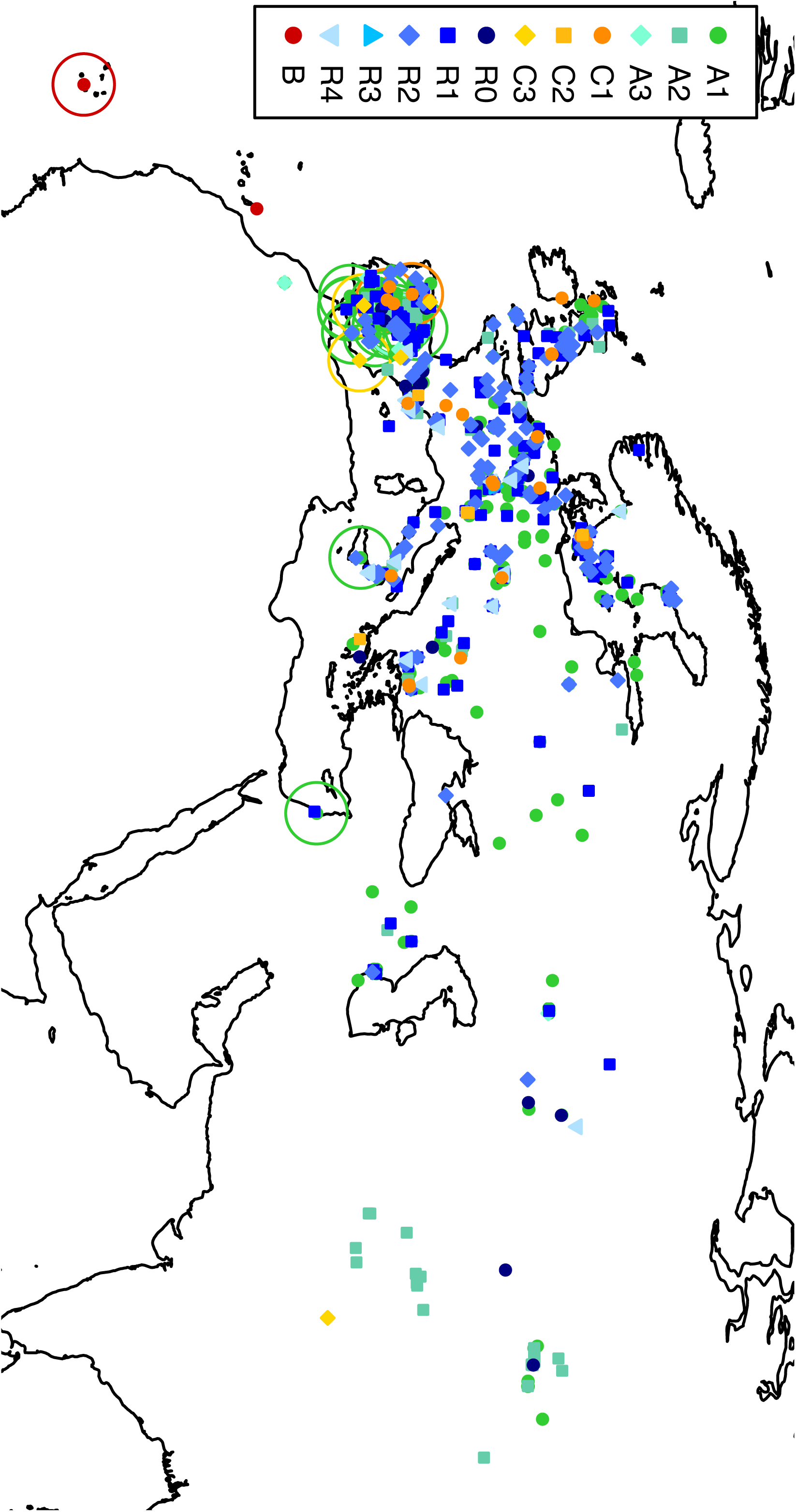
Geographical distribution of the main haplogroups identified. Each accession is represented by a color-coded dot on the map. ‘Relict’ accessions are circled with a circle in the background, colored according to the S-haplotype assignment.

Genome-wide patterns of divergence identified several groups of ‘Relict’ accessions that directly derive from glacial refugia, with very limited amount of admixture since then (The 1001 Genomes Consortium, 2016). If the breakdown of SI had occurred within these refugia, then S-haplotypes would have become differentially fixed locally and the ‘Relict’ accessions would be monomorphic, each with a different haplogroup at the S-locus. This prediction may hold true for the glacial refugia in offshore Africa where haplogroup B was detected, but since the relict accessions from Iberia contained both haplogroups A and C, it seems likely that the breakdown of SI in *A*. *thaliana* occurred prior to the last Ice Age, and that the dramatic reduction of the number of S-haplogroups (as compared to self-incompatible species) associated with the breakdown of SI had already occurred when the glacial refugia were established.

### Frequent recombinant haplotypes and their origins

Recombination between S-locus haplotypes is believed to be suppressed because of extreme sequence divergence, and to be selectively disfavored because of functional constraints (Goubet et al. 2012). Thus, the high frequency of inter-haplogroup recombinants, with as many as 51.5% of accessions aligning substantially to both references of haplogroups A and C, is very striking.

To understand the origin of the recombinants, we inspected the recombination breakpoints in detail. A single recombinant haplotype sequence had previously been reported (C24; Sherman-Broyles et al. 2007), and an independent recombination event has also been suggested (Kas-2, Boggs et al. 2009). These studies suggested that the C24 recombinant haplotype could have been formed via a partially duplicated flanking gene, *ARK3* (called *ΔARK3*), which could have acted as the recombination breakpoint (Sherman-Broyles et al. 2007, Dwyer et al. 2013). Our mapping data are also consistent with the scenario that *ΔARK3* in one parental C haplotype recombined heterologously with the original *ARK3* from an A-haplotype in heterozygous A/C plants (Fig. S5), as recombinants have substantial coverage that extends up to this point. We used long-range PCR to compare the flanking sequences of *ΔARK3* in a set of eleven R1, R2, R3 or R4 haplotypes (hence spanning across the A-C boundary) and found that they were almost identical to those in C24, confirming that their *ΔARK3* were inserted at orthologous positions and therefore derive from a single duplication event.

To identify the precise location of the breakpoints, we performed Sanger sequencing of the recombination breakpoint proposed in C24 (Sherman-Broyles et al. 2007). In line with Tsuchimatsu et al. (2010), we found that the vast majority of the observed polymorphisms could be classified into one of two divergent haplogroups of *ARK3,* which enabled us very precisely to pinpoint the position of the recombination breakpoint, to within a few nucleotides. Strikingly, we found at least seven distinct breakpoints (Fig. 6). Although gene conversion from the *ARK3* gene cannot be excluded, these results suggest that the distinct breakpoints represent as many independent recombination events, and that once the *ΔARK3* sequence was in place, haplogroups A and C repeatedly recombined.

**Figure 6.**
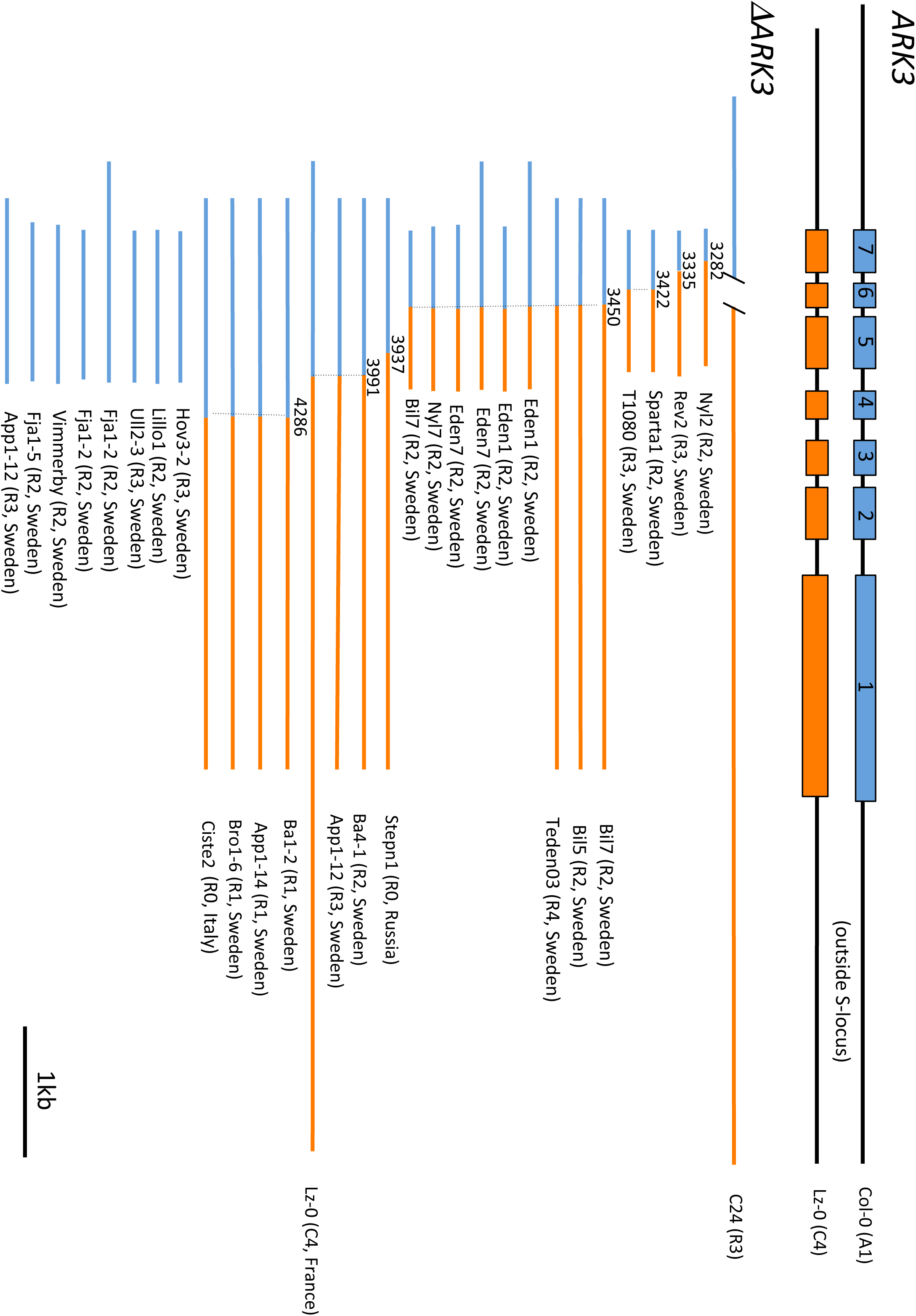
Sanger resequencing identifies several recombination breakpoints at the *ΔARK3* region. Blue and orange colors identify the divergent *ARK3* haplotypes. Some accessions were sequenced from multiple pairs PCR primers and in such case they were represented several times. We note that the deletion encompassing exon 6 of *ΔARK3,* which was reported previously in the C24 accession, was not found in other accessions, suggesting its uniqueness to C24. We confirmed that it indeed exists in the C24 accession by Sanger sequencing.

### Lack of recombinants in relict accessions

Despite the overall high frequency of recombinants, we note that not a single of the 22 ‘Relict’ accessions carried a recombinant haplogroup, a significant underrepresentation (chi square = 12.65, df = 1, p=0.0005). Hence, recombinants are specific to the accessions derived from the ‘invasive’ refugium that recolonized most of Eurasia and recombination seems to have taken place on the specific genetic background carrying *ΔARK3* sequence. It is therefore tempting to speculate that recombination between distinct S-haplogroups played a direct role in the loss of SI, as suggested by Sherman-Broyles et al. (2007). To test this, we examined the *SCR* sequence and found that all the haplotypes containing the *SCR-A* region (A1, A3, R0 and R1 haplotypes) harbor the 213-bp inversion in *SCR-A,* which was previously suggested to be the mutation responsible for the loss of SI in haplogroup A (Tsuchimatsu et al. 2010), and also share most of the derived mutations that potentially deactivate *SRK.* This strongly suggests that the haplotypes that gave rise to the recombinants were already nonfunctional.

## Discussion

### Geographic distribution of S-haplogroups, post-glacial colonization, and multiple origins of self-compatibility

Although the shift to selfing is common in flowering plants, the scenario under which this occurs varies across species (Vekemans et al. 2014). *Capsella rubella,* for example, has undergone a global fixation of a single non-functional S-allele associated with speciation (Guo et al. 2009). In contrast, in *A*. *lyrata* non-functional alleles segregate in some populations along with functional alleles, and in *Leavenworthia alabamica* different selfing populations have fixed different non-functional alleles, so that no species-wide fixation of the non-functional alleles is observed in either species (Mable et al. 2005, 2016; Busch et al 2011).

In *A*. *thaliana* we find yet another pattern, combining species-wide loss of SI with the maintenance of at least three non-functional S-haplogroups (given the large sample analyzed here, from a broadly distributed set of populations, it is unlikely that additional haplotypes, if they exist, have substantial frequencies). Given that the number of S-haplotypes that segregated in the ancestral outcrossing species was very large, this entails a major reduction in the number of SI alleles (Fig. 7). Still, the existence of more than one non-functional haplotype is puzzling, and raises the question why the first disrupted haplotype did not become fixed species-wide if the conditions for invasion were met. A first possibility is that different selfing genotypes became fixed in separate glacial refugia. This might be true for haplotype B, which we found to be restricted to offshore Africa. However, the widespread geographical distribution of both haplogroups A and C shows that their putative fixation did not occur within specific refugia from the last glaciation. We even found that both haplotypes segregate among relict accessions from the single Iberian refugium, demonstrating that they were already co-occurring when the refugium was formed (Fig. 7). Hence, local fixations of alternative S-haplotypes in refugia, if they happened, must have taken place during an earlier glacial cycle, followed by secondary admixture during a previous interglacial period.

**Figure 7.**
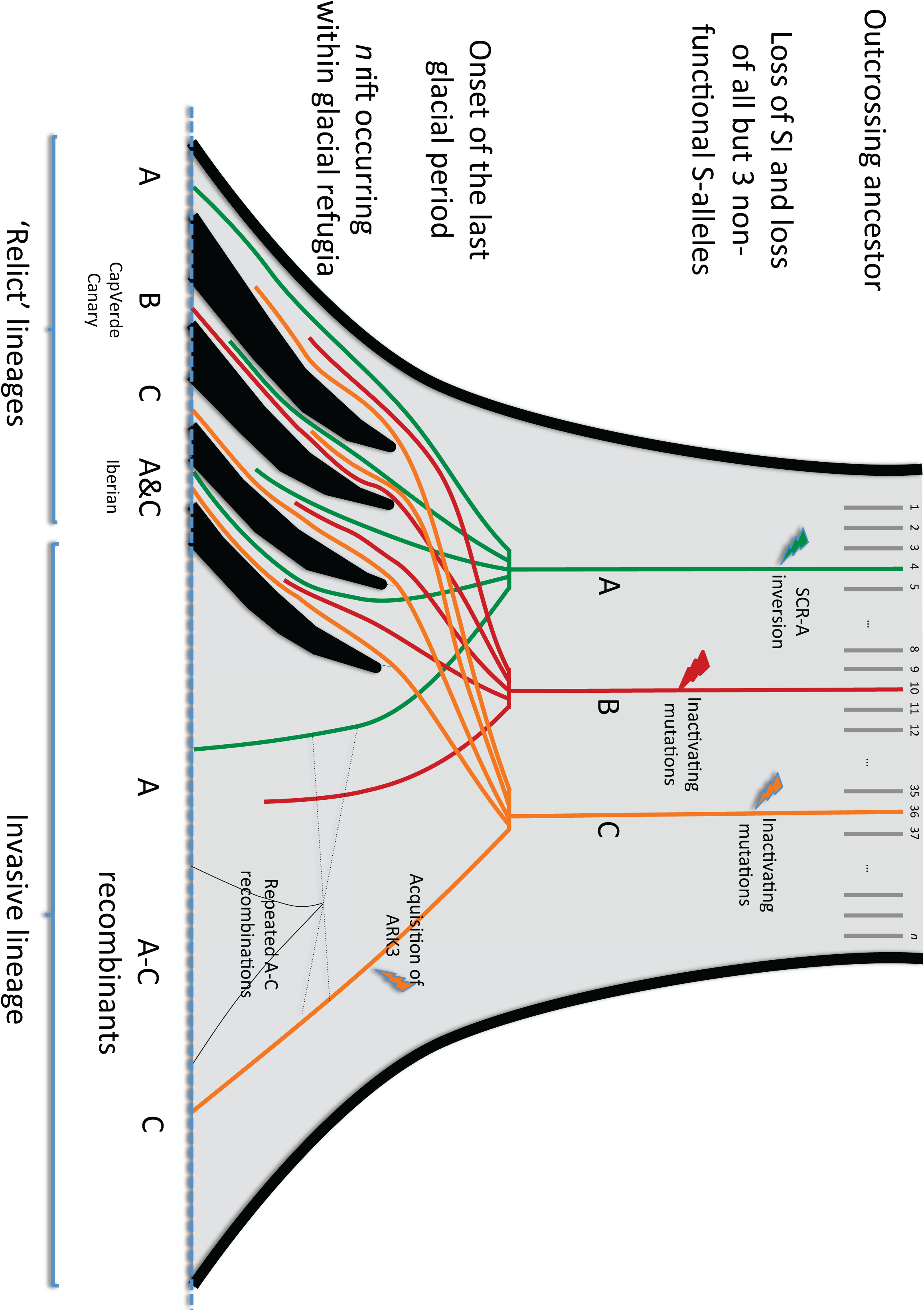
Scenario for the multiple loss of self-incompatibility, subsequent decay of the S-locus and distribution of S-haplotypes in different glacial refugia. The number of functional S-haplotypes was initially very large in the outcrossing ancestor, while only three S-haplogroups currently segregate, each containing specific inactivating mutations. As glacial refugia became isolated, genetic drift led to differential fixation of S-haplogroups in the different refugia, while others remained polymorphic (as in the Iberian refugium). The lineage from the refugia that became invasive thoughout Eurasia specifically acquired a local duplication of the flanking gene *ARK3* on haplogroup C that allowed for repeated recombination events among copies of the different S-haplogroups that were locally segregating.

A second possibility is that the species-wide fixation process was slow, providing enough time for the emergence of several non-functional haplotypes. Theory predicts that non-functional haplotypes may stably segregate in a self-incompatible population along with functional haplotypes, specifically when inbreeding depression is high and the proportion of self-pollen deposited on pistils is low to intermediate (Charlesworth and Charlesworth 1979; Uyenoyama et al. 2001; Gervais et al. 2011). If so, multiple non-functional haplotypes could have sequentially appeared and been maintained within the species. Because all three haplotypes carry inactivating mutations, this polymorphism could have been effectively neutral and may be on its way to slow fixation by genetic drift. Population structure, which is currently strong in *A*. *thaliana* (Nordborg et al. 2005), would further slow down the species-wide fixation of a particular non-functional haplotype. In addition, the discovery of a fixed deletion in *A*. *thaliana* of a putative gene of the signaling cascade of SI (Indriolo et al. 2012, but see Nasrallah & Nasrallah 2014) suggests the possibility that the ancestral *A*. *thaliana* lineage experienced partial self-compatibility. This could have contributed to the reduction of the selective forces promoting non-functional S-haplotypes, hence slowing down their fixation.

### Extensive recombination at the S-locus in a selfing species

Recombination between functional SI haplotypes is apparently suppressed in outcrossing species, but it is currently unclear whether recombination suppression is mainly due to selective constraints or to the major structural polymorphism preventing proper chromosomal pairing during meiosis (Castric et al. 2010). The overall frequency of recombinant haplotypes we observed is much higher than previous estimates (22.86%, Sherman-Broyles et al. 2007), possibly because previous genotyping was based on detecting small gene fragments that we found to have been subsequently deleted in specific haplotypes (Fig. S2). With the whole S-locus sequence, we have much greater power to detect recombination events even after subsequent deletions have occurred.

Our data show that in self-compatible *A*. *thaliana*, where functional constraints are absent, recombination has occurred repeatedly between a pair of highly diverged haplogroups. Hence, the extreme level of structural polymorphism among haplogroups does not seem to prevent recombination. We note, however, that the signatures of inter-haplogroup recombination indicate that it is caused by unequal crossing-over involving a duplicated copy of the flanking gene *ARK3* (Fig S5). Interestingly, *ARK3* seems to be prone to duplications even in functional SI haplotypes, as distinct *ΔARK3-like* sequences have been found in multiple haplotypes in *A*. *lyrata* and *A*. *halleri* (Guo et al. 2011, Goubet et al. 2012), and a polymorphism survey at *ARK3* in *A*. *lyrata* has reported the frequent occurrence of multiple copies (Hagenblad et al. 2006), a property that could facilitate rearrangements of the S-locus.

### Structural rearrangements as secondary decay

An important challenge in the studies on the loss of SI has been to distinguish primary inactivating mutations causing loss of SI from subsequent decay by further gene-disruptive mutations and structural rearrangements (Igic et al. 2008, Busch and Schoen 2008, Tsuchimatsu et al. 2010, Boggs et al. 2009). Sherman-Broyles et al. (2007), who first reported the extensive structural rearrangements at the S-locus region of C24, proposed two scenarios for the loss of SI in the recombinant haplotypes: (1) recombination between the haplogroups A and C in a heterozygous self-incompatible individual causing loss of SI, followed by subsequent restructuring, or (2) inactivation of the haplogroups A and C by independent rearrangements and deletions followed by recombination in a self-fertile heterozygous individual. We found that all recombinant haplotypes containing the *SCR-A* region had the 213-bp inversion in *SCR-A,* which was reported to be responsible for the loss of SI within haplogroup A (Tsuchimatsu et al. 2010). This result suggests that pseudogenization of *SCR-A* preceded the recombination events as well as the large deletions. In addition, our results suggest that the parental haplotype carrying haplogroup C was also non-functional due to the loss of the *SCR* promoter region, such that we have high confidence that the SI system of the A/C heterozygote was already deactivated. Finally, the fact that none of the relict accessions had any of the large-scale rearrangements we identified, strongly suggests that those rearrangements happened in the very recent past, possibly post-glacially, and were specific to the invasive “weedy” lineage that recolonized most of Eurasia (Fig. 7). At any rate, it is now clear that the independent recombination events and their associated structural rearrangements represent truly secondary decay that occurred after the loss of SI of both parental haplotypes.

## Materials and Methods

### Isolation of a genomic fragment containing the A. thaliana C haplogroup

Isolation of a genomic fragment containing the *A*. *thaliana* C haplogroup was performed according to Goubet et al. (2012). Briefly, high molecular weight DNA was prepared from young leaves of the Ita-0 accession and a BAC library was constructed. The library was screened on nylon filters using radiolabelled probes designed from the flanking genes *ARK3* and *PUB8*. Positive BAC clones detected by hybridization were validated by PCR amplification using the primer pairs used for probes synthesis, and visualisation of PCR products after agarose gel electrophoresis. The BAC clone was sequenced at Genoscope using a 454 multiplexing technology on the Titanium sequencer version (www.roche.com). *De novo* assembly was performed by Newbler (www.roche.com) and the sequence was obtained in five contigs. Only contigs representing the extremities of the BAC were initially oriented. Long-range PCR was then employed to order and orientate the rest of the contigs (Table S1). Genes and transposable elements annotation was performed according to Goubet et al. (2012). Because the Ah36 sequence published in Durand et al. (2014) contains an unusual duplicated copy of *SCR*, we resequenced this BAC clone using the PACBIO platform (Genbank accession number XXXX) and confirmed this structural feature.

### Expression analysis

Total RNAs were extracted from stigma (S) and anthers (A) (stage 12 according to Smyth et al. 1990) of *A*. *lyrata* S14 haplotype and *A*. *thaliana* Ita-0 plants following the manufacturer’s recommendations (Picopure kit; Arcturus) except that a second DNase treatment (RNase-free Dnase, Qiagen) was added. The complementary DNA (cDNA) was synthesized with the Revert Aid M-MuLV reverse transcription (Fermentas) using (N)6 random primers and 260ng of total RNAs. Reverse transcription (RT)-PCR with 25 cycles for *SRK* amplification or 35 cycles for *SCR* amplification was performed with primer pairs located within the exon 1 for *SRK14*, surrounding the intron 1 for *SRKIta-0* and on the ATG and the stop codons for *SCR.* The constitutively expressed gene *ACT8* was used as a control to normalize loading of RT-PCR products. Amplification using primers located within a non-expressed region (intergenic region between At1g49230 and At1g49240 (ACT8)) confirmed the absence of DNA contamination (data not shown). Primer pairs for *SRKIta-0* and *SCRIta-0* were tested using, as template, BAC DNA (BAC) containing the S-locus of Ita-0. The PCR products amplified correspond to the genomic fragments including introns (1229 and 1288 bp respectively).

### Analysis of the S-locus region in accessions from the 1,001 genomes project

We used the paired-end reads from whole genome sequencing of 1,083 accessions from the 1,001 genomes project. Briefly, each accession was sequenced using Illumina sequencing with paired-end reads of 76 bp. Because of the large sequence divergence among haplotypes, the S-locus can typically not be assembled using standard procedures (*i.e.* mapping on a single genomic reference, Col-0). Hence, for each of these accessions, three new alignments of the S-locus were performed using the Burrows-Wheeler Aligner (Li and Durbin 2009), taking in turn each known sequence (Col-0 for haplogroup A, Cvi-0 for haplogroup B, Ita-0 for haplogroup C) as a reference.

We then plotted for each accession the proportion of nucleotide positions showing non-zero coverage (>1 reads) on references of haplogroups A, B and C. We masked transposable elements (TE) for this analysis, as the extremely high coverage on TEs would be mainly due to non-specific reads and thus confound the results. This analysis revealed a discrete set of coverage patterns, suggesting a limited number of S-locus types among accessions. Based on a representative subset of accessions for each main type, we then searched manually for fragments of the reference sequences that distinguished among these main types. While most fragments varied in a highly coordinated manner, a 12kb fragment of Ita-0 (represented by a dashed motif on Fig. 1 and S2) was not specific to accessions defined as haplotype C or R by all other fragments. This fragment also varied in a presence/absence manner (data not shown) and had substantial coverage in about 45% of all accessions but we could not confirm contiguity between this fragment and its flanking sequences on the C-reference by any paired-end sequencing read straddling these positions. To further examine the origin of this fragment, we extracted paired end reads for which one read mapped at one extremity of the fragment and the other was unmapped on the references. We then blasted the unmapped reads on the complete *A*. *thaliana* genome and found for the two extremities a strong homology with one particular position on chromosome 2 (around position 1,825,600). This observation suggests that this 12kb fragment segregates on chromosome 2 of 45% of all accessions and has been specifically inserted in the S-locus of the Ita-0 accession. Consequently, we decided to ignore its pattern of presence/absence in the analyzed accessions. Overall, this analysis allowed us to distinguish a total of 12 different types that varied by their pattern of presence/absence of the different fragments. We used this set of fragments to automatically classify the rest of the accessions. A fragment was considered to be present if the proportion of sites with non-zero coverage was more than the threshold value defined for each fragment, and otherwise considered to be absent.

Based on the distribution of the median pairwise distance to all other accessions computed from all SNPs (The 1001 Genomes Consortium 2016), we chose to conservatively consider as relict accessions those with a median distance above 0.00356, corresponding to a gap in the distribution. Application of this strict threshold resulted in 22 relict accessions. According to The 1001 Genomes Consortium (2016), the majority of these relicts were from the Iberian peninsula (*n*=19) and are considered as descendants from a single glacial refugium, with one accession each representing relicts from Sicily (*n*=1), Lebanon (*n*=1) and the Cape-Verde island (*n*=1).

### Targeted resequencing across the recombination breakpoint

We obtained sequences around the recombination breakpoints and the deletion border by Sanger sequencing. DNA was extracted by Macherey-Nagel plant DNA kit. GoTaq (Promega) was used for PCR. PCR products were purified by using Agencourt AMPure XP (Beckman Coulter) and sequenced by Applied Biosystems 3130xL capillary sequencer. For primers used, see Table S1. Sequences were inspected by eye, assembled and aligned by using the CLC Main Workbench. For long-range PCR fragments, internal primers were designed to tile the fragments entirely.

## Acknowledgements

We thank Envel Kerdaffrec for experimental support and Deborah Charlesworth for comments on the manuscript. PG was supported by a PhD fellowship from CNRS. EMBO long-term fellowship, JSPS postdoctoral fellowship for research abroad, and JSPS KAKENHI (15K18583) to TT are gratefully acknowledged. VC benefited from a Fulbright fellowship. BAC library construction was performed in the frame of the ATIP+ program from CNRS to XV. BAC sequencing was performed by Genoscope in the frame of Project 2006#13 to VC and XV. Financial support from French Agence Nationale de la Recherche (ANR-11-BSV7-013-03, ANR-11-JSV7-008-01) and European Research Council (NOVEL project, grant #648321) is also gratefully acknowledged. This paper is dedicated to the memory of one of its co-first authors Pauline Goubet, who passed away before this work was finalized.

## Supplementary Material

**Figure S1.** Phylogenetic tree of *SRK-C* amino acid sequences from a subset of haplogroup C and R accessions (R0 haplotype) of *A*. *thaliana* together with sequences from the outcrossing relatives *A. lyrata (AlSRK36)* and *A*. *halleri* (AhSRK36). The phylogeny was obtained with the neighbor-joining algorithm in MEGA5 (Tamura et al. 2011) with 1,000 bootstrap replicates. Only bootstrap values above 50% are reported.

**Figure S2.** Patterns of presence/absence of the different fragments detected in the main haplogroups identified. Fragments used for haplogroup assignment are represented as red segments.

**Figure S3.** Distribution of coverage across the 1083 accessions for the different fragments identified on Fig. S2. The vertical red line corresponds to the arbitrary threshold set to define a fragment as being present or absent.

**Figure S4.** Geographical distribution of identified haplotypes plotted for each haplogroup (A, B, C and the recombinants).

**Figure S5.** A proposed scenario for the putative recombination event between haplogroups A and C. The first step is the duplication of a portion of the flanking gene *ARK3* providing the substrate of sequence similarity allowing heterologous recombination.

## REFERENCES

Abbott RJ, Gomes MF (1989) Population genetic structure and outcrossing rate of *Arabidopsis thaliana* (L.) Heynh. Heredity 62: 411–418.

Arabidopsis Genome Initiative (2000) Analysis of the genome sequence of the flowering plant *Arabidopsis thaliana*. Nature 408: 796–815.

Barrett SCH (2002) The evolution of plant sexual diversity. Nat Rev Genet 3: 274–284.

Bechsgaard JS, Castric V, Charlesworth D, Vekemans X, Schierup MH (2006) The transition to self-compatibility in *Arabidopsis thaliana* and evolution within *S*-haplotypes over 10 Myr. Mol Biol Evol 23: 1741–1750.

Boggs NA, Nasrallah JB, Nasrallah ME (2009) Independent *S*-locus mutations caused self-fertility in *Arabidopsis thaliana*. PLoS Genet 5: e1000426.

Bomblies K, Yant L, Laitinen RA, Kim S-T, Hollister JD, et al. (2010) Local-scale patterns of genetic variability, outcrossing, and spatial structure in natural stands of *Arabidopsis thaliana*. PLoS Genet 6: e1000890.

Busch JW, Joly S, Schoen DJ (2011) Demographic signatures accompanying the evolution of selfing in *Leavenworthia alabamica*. Mol Biol Evol 28: 1717–1729.

Busch JW, Schoen DJ (2008) The evolution of self-incompatibility when mates are limiting. Trends Plant Sci 13: 128–136.

Cao J, Schneeberger K, Ossowski S, Günther T, Bender S, et al. (2011) Whole-genome sequencing of multiple *Arabidopsis thaliana* populations. Nat Genet 43: 956–963.

Castric V, Bechsgaard JS, Grenier S, Noureddine R, Schierup MH, et al. (2010) Molecular evolution within and between self-incompatibility specificities. Mol Biol Evol 27: 11–20.

Charlesworth D, Charlesworth B (1979) The evolution and breakdown of S-allele systems. Heredity 43: 41–55.

Darwin C (1876) The effects of cross and self fertilisation in the vegetable kingdom. London: J. Murray. 487 p.

de Nettancourt D (2001) Incompatibility and incongruity in wild and cultivated plants, 2nd edn. Berlin: Springer. 322 p.

Durand E, Méheust R, Soucaze M, Goubet PM, Gallina S et al. (2014) Dominance hierarchy arising from the evolution of a complex small RNA regulatory network. Science 346: 1200–1205.

Dwyer KG, Berger MT, Ahmed R, Hritzo MK, McCulloch AA, et al. (2013) Molecular characterization and evolution of self-incompatibility genes in *Arabidopsis thaliana:* the case of the *Sc* haplotype. Genetics 193: 985–994.

Fisher RA (1941) Average excess and average effect of a gene substitution. Ann Eugen 11: 53–63.

Gan X, Stegle O, Behr J, Steffen JG, Drewe P, et al. (2011) Multiple reference genomes and transcriptomes for *Arabidopsis thaliana*. Nature 477: 419–423.

Gervais CE, Castric V, Ressayre A, Billiard S. 2011. Origin and diversification dynamics of self-incompatibility haplotypes. Genetics 188 (3): 625–636.

Gervais C, Awad DA, Roze D, Castric V, Billiard S (2014) Genetic architecture of inbreeding depression and the maintenance of gametophytic self-incompatibility. Evolution 68: 3317–3324.

Goring DR, Indriolo E, Samuel MA (2014) The *ARC1* E3 ligase promotes a strong and stable self-incompatibility response in *Arabidopsis* species: response to the Nasrallah and Nasrallah commentary. Plant Cell 26: 3842–3846.

Goubet PM, Bergès H, Bellec A, Prat E, Helmstetter N, et al. (2012) Contrasted Patterns of Molecular Evolution in Dominant and Recessive Self-Incompatibility Haplotypes in *Arabidopsis*. PLoS Genet 8: e1002495.

Guo YL, Zhao X, Lanz C, Weigel D (2011) Evolution of the *S*-locus region in *Arabidopsis* relatives. Plant Physiol 157: 937–946.

Hagenblad J, Bechsgaard J, Charlesworth D (2006) Linkage disequilibrium between incompatibility locus region genes in the plant *Arabidopsis lyrata*. Genetics 173: 1057–1073.

Igic B, Lande R, Kohn JR (2008) Loss of self-incompatibility and its evolutionary consequences. Int J Plant Sci 169: 93–104.

Indriolo E, Safavian D, Goring DR (2014) The *ARC1* E3 ligase promotes two different self-pollen avoidance traits in *Arabidopsis*. Plant Cell 26: 1525–1543.

Indriolo E, Tharmapalan P, Wright SI, Goring DR (2012) The *ARC1* E3 ligase gene is frequently deleted in self-compatible Brassicaceae species and has a conserved role in *Arabidopsis lyrata* self-pollen rejection. Plant Cell 24: 4607–4620.

Kitashiba H, Liu P, Nishio T, Nasrallah JB, Nasrallah ME (2011) Functional test of *Brassica* self-incompatibility modifiers in *Arabidopsis thaliana*. Proc Natl Acad Sci U S A 108: 18173–18178.

Kusaba M, Dwyer K, Hendershot J, Vrebalov J, Nasrallah JB, et al. (2001) Self-incompatibility in the genus *Arabidopsis*: Characterization of the *S-*locus in the outcrossing *A. lyrata* and its autogamous relative *A. thaliana*. Plant Cell 13: 627–643.

Li H, Durbin R (2009) Fast and accurate short read alignment with Burrows-Wheeler transform. Bioinformatics 25: 1754–1760.

Liu P, Sherman-Broyles S, Nasrallah ME, Nasrallah JB (2007) A cryptic modifier causing transient self-incompatibility in *Arabidopsis thaliana*. Curr Biol 17: 734–740.

Llaurens V, Billiard S, Leducq JB, Castric V, Klein EK, et al. (2008) Does frequency-dependent selection with complex dominance interactions accurately predict allelic frequencies at the self-incompatibility locus in *Arabidopsis halleri*? Evolution 62: 2545–2557.

Long Q, Rabanal FA, Meng D, Huber CD, Farlow A, et al. (2013) Massive genomic variation and strong selection in *Arabidopsis thaliana* lines from Sweden. Nat Genet 45: 884–890.

Mable BK, Schierup MH, Charlesworth D (2003) Estimating the number, frequency, and dominance of S-alleles in a natural population of *Arabidopsis lyrata* (Brassicaceae) with sporophytic control of self-incompatibility. Heredity 90: 422–431.

Mable B, Robertson A, Dart S, Di Berardo C, Witham L (2005) Breakdown of self-incompatibility in the perennial *Arabidopsis lyrata* (Brassicaceae) and its genetic consequences. Evolution 59: 1437–1448.

Mable BK, Hagmann J, Kim ST, Adam A, Kilbride E, Weigel D, Stift M (2016) What causes mating system shifts in plants? Arabidopsis lyrata as a case study. Heredity. in press. doi:10.1038/hdy.2016.99

Murase K, Shiba H, Iwano M, Che F-S, Watanabe M, et al. (2004) A membrane-anchored protein kinase involved in *Brassica* self-incompatibility signaling. Science 303: 1516–1519.

Nasrallah JB, Nasrallah ME (2014) Robust self-incompatibility in the absence of a functional *ARC1* gene in *Arabidopsis thaliana*. Plant Cell 26: 3838–3841.

Nasrallah ME, Liu P, Nasrallah JB (2002) Generation of self-incompatible *Arabidopsis thaliana* by transfer of two *S* locus genes from *A. lyrata*. Science 297: 247–249.

Nasrallah ME, Liu P, Sherman-Broyles S, Boggs NA, Nasrallah JB (2004) Natural variation in expression of self-incompatibility in *Arabidopsis thaliana:* implications for the evolution of selfing. Proc Natl Acad Sci U S A 101: 16070–16074.

Nordborg M, Hu TT, Ishino Y, Jhaveri J, Toomajian C, et al. (2005) The pattern of polymorphism in *Arabidopsis thaliana*. PLoS Biol 3: e196.

Platt A, Horton M, Huang YS, Li Y, Anastasio AE, et al. (2010) The scale of population structure in *Arabidopsis thaliana*. PLoS Genet 6: e1000843.

Samuel MA, Chong YT, Haasen KE, Aldea-Brydges MG, Stone SL, et al. (2009) Cellular pathways regulating responses to compatible and self-incompatible pollen in *Brassica* and *Arabidopsis* stigmas intersect at Exo70A1, a putative component of the exocyst complex. Plant Cell 21: 2655–2671.

Sherman-Broyles S, Boggs N, Farkas A, Liu P, Vrebalov J, et al. (2007) *S* locus genes and the evolution of self-fertility in *Arabidopsis thaliana*. Plant Cell 19: 94–106.

Shimizu KK, Shimizu-Inatsugi R, Tsuchimatsu T, Purugganan MD (2008) Independent origins of self-compatibility in *Arabidopsis thaliana*. Mol Ecol 17: 704–714.

Shimizu KK, Tsuchimatsu T. 2015. Evolution of Selfing: Recurrent Patterns in Molecular Adaptation. Annual Review of Ecology, Evolution, and Systematics 46: 593–622.

Smyth DR, Bowman JL, Meyerowitz EM (1990) Early flower development in *Arabidopsis*. Plant Cell 2: 755–767.

Stebbins GL (1974) Flowering plants: Evolution above the species level. Cambridge: Belknap Press. 399 p.

Stone S, Anderson E, Mullen R, Goring D (2003) ARC1 is an E3 ubiquitin ligase and promotes the ubiquitination of proteins during the rejection of self-incompatible *Brassica* pollen. Plant Cell 15: 885–898.

Takayama S, Isogai A (2005) Self-incompatibility in plants. Annu Rev Plant Biol 56: 467–489.

Tamura K, Peterson D, Peterson N, Stecher G, Nei M, et al. (2011) MEGA5: Molecular evolutionary genetics analysis using maximum likelihood, evolutionary distance, and maximum parsimony methods. Mol Biol Evol 28: 2731–2739.

Tang C, Toomajian C, Sherman-Broyles S, Plagnol V, Guo YL, et al. (2007) The evolution of selfing in *Arabidopsis thaliana*. Science 317: 1070–1072.

The 1001 Genomes Consortium (2016) 1,135 genomes reveal the global pattern of polymorphism in *Arabidopsis thaliana*. Cell 166: 481–491.

Tsuchimatsu T, Suwabe K, Shimizu-Inatsugi R, Isokawa S, Pavlidis P, et al. (2010) Evolution of self-compatibility in *Arabidopsis* by a mutation in the male specificity gene. Nature 464: 1342–1346.

Uyenoyama MK, Zhang Y, Newbigin E (2001) On the origin of self-incompatibility haplotypes: Transition through self-compatible intermediates. Genetics 157: 1805–1817.

Vekemans X, Poux C, Goubet PM, Castric V (2014) The evolution of selfing from outcrossing ancestors in Brassicaceae: what have we learned from variation at the S-locus? J Evol Biol 27: 1372–1385.

